# Molecular activity mediates the composition and assembly of dissolved organic matter in lake sediments

**DOI:** 10.1101/2024.09.17.613390

**Authors:** Shuailong Wen, Ang Hu, Francisco Dini-Andreote, Lei Han, Shuyu Jiang, Kyoung-Soon Jang, Jianjun Wang

**Affiliations:** Key Laboratory of Lake and Watershed Science for Water Security, Nanjing Institute of Geography and Limnology, Chinese Academy of Sciences, Nanjing 210008, China; State Key Laboratory of Lake Science and Environment, Nanjing Institute of Geography and Limnology, Chinese Academy of Sciences, Nanjing 210008, China; Department of Plant Science & Huck Institutes of the Life Sciences, The Pennsylvania State University, University Park, PA, USA; The One Health Microbiome Center, Huck Institutes of the Life Sciences, The Pennsylvania State University, University Park, PA, USA; Bio-Chemical Analysis Team, Korea Basic Science Institute, Cheongju, Republic of Korea

**Author notes:** Corresponding Author: Kyoung-Soon Jang, Jianjun Wang.

**Keywords:** molecular activity, potential biochemical transformations, dissolved organic matter, lake sediments, assembly processes, bacterial communities

## Abstract

Lake sediments are hotspots for carbon transformation and burial, where dissolved organic matter (DOM) interacts with microorganisms to regulate global carbon cycling. The potential for individual DOM molecules to undergo biochemical transformations, i.e., their activity, is a critical molecular trait affecting DOM turnover in environment. However, the composition of sediment DOM and how its assembly mechanisms are related to molecular activity remains poorly understood. Here, 63 freshwater sediments were collected from tropical to cold temperate climatic zones in China. We explored the molecular composition and assembly of sediment DOM and the underlying mechanisms driven by climate, physicochemical factors, and microbes along the molecular activity gradient. Sediment DOM was dominated by lipid- (34.8%) and lignin-like compounds (33.01%), and the latter were enriched as molecular activity of DOM increased. Besides, DOM composed of more active molecules had greater compositional similarity across different climatic zones, and was inclined to assemble deterministically. This was supported by the fact that as potential transformations of molecular assemblages increased from 0.4 to 14, the assembly of these molecules was structured by a shift from stochastic to deterministic processes, with the latter accounting for ≥ 75% thereafter. Overall, the molecular assemblage was primarily structured by physicochemical factors, including sediment total organic carbon and electrical conductivity. As molecular activity increased, however, assemblage was increasingly affected by climate and bacterial communities, consistent with the enhanced complexity of bacterial-molecular networks. Collectively, our study highlights that the intrinsic activity of DOM molecules determines their fate through distinct biotic and abiotic mechanisms.

## Introduction

As the most dynamic component for carbon cycling in waters and sediments, dissolved organic matter (DOM) exerts profound effects on the functionality of lakes and downstream aquatic ecosystems (Cui et al., 2024; Tanentzap and Fonvielle, 2024). These effects include the promotion of eutrophication (Liu et al., 2022), the potential alteration of contaminant toxicity (Gonsior et al., 2019), and the modulation of biogeochemical cycles (e.g., nitrogen, carbon, sulfur) via shifts in microbial metabolisms (Brailsford et al., 2021; Chen et al., 2022). Therefore, DOM composition – which refers to the chemodiversity and structural properties of the organic compounds – has been broadly explored in overlying water bodies (e.g., rivers, lakes, and wetlands) (Lapierre et al., 2013; Tanentzap and Fonvielle, 2024). However, knowledge on the composition of DOM in sediments, which represent one of the largest reservoirs of active carbon in freshwater ecosystems (Catalán et al., 2016), remains still largely elusive. As such, understanding the underlying mechanisms structuring the composition of DOM in freshwaters currently represents a fundamental knowledge gap.

DOM composition in freshwaters is determined by a complex interplay among several forces, including production, degradation, transport, and transformation of organic molecules (Danczak et al., 2020). Most interestingly, these interplays can occur through a balance of deterministic and stochastic mechanisms (Kajan et al., 2023). Analogous to the concept of ecological community assembly (Ning et al., 2020), deterministic processes are associated with the selective production, transformation, or loss of specific molecules through the interactions among biotic and abiotic factors, also known as environmental filtering (Hu et al., 2022b). For instance, microorganisms preferentially degrade lipid- and protein-like molecules, causing an enrichment of polyphenolic and highly unsaturated recalcitrant compounds in lakes (Catalán et al., 2024). Besides, DOM composition could be determined by stochastic processes, which involve random events (in ecology, dispersal and ecological drift) – here, the hydrological transport and vectorial movement of molecules and specific molecules’ lifetime (Hu et al., 2022b; Hubbell, 2001).

Organic molecules undergo numerous biochemical transformations, with consequences for the composition and assembly of DOM that are highly related to molecular reactivity and activity (Kajan et al., 2023; Ryan et al., 2024). Unlike the well-studied reactivity, which describes a molecule’s ability to be decomposed, molecular activity – indicating the count of potential transformations a molecule can undergo – has rarely been explored in relation to DOM composition and assembly across divergent systems (Hu et al., 2022b; Lau and Del Giorgio, 2020). For instance, the main transformation pathways related to molecular reactivity, such as biological reactions (Mostovaya et al., 2016), sunlight-induced photochemical reactions (Grasset et al., 2024), and immobilization by flocculation (Eusterhues et al., 2011), have been extensively explored for revealing DOM dynamics in terrestrial and freshwater habitats (Berggren et al., 2022). However, the effects of molecular activity on DOM composition and assembly are poorly understood. Recently, it has been shown that mass differences between paired molecules resolved by high-resolution mass spectrometry could be used to infer potential transformations of molecules (Danczak et al., 2020; Kajan et al., 2023). These potential transformations provide valuable insights into the molecular activity associated with DOM dynamics and assembly, despite it does not directly inform when or where a particular transformation occurs (Li et al., 2023a; Stegen et al., 2022). Here, we hypothesize that DOM would be assembled more deterministically as molecular activity increases. This is supported by the notion that the molecular assemblages within high potential transformations tend to be more closely associated with microbial communities (Danczak et al., 2020; Li et al., 2023a). Besides, we hypothesize that the underlying drivers mediating DOM composition to vary along the gradient of molecular activity, due to direct effects of microbes and indirect effects of physicochemical properties on the transformation of sediment DOM molecules.

To verify the above hypotheses, we studied variations in the composition and assembly mechanisms of continental-scale sediment DOM along a gradient of molecular activity represented by potential biochemical transformations. We collected 63 freshwater sediments from tropical to cold temperate climatic zones in China that spanning 31 degrees of latitude, and analyzed their DOM composition by ultra-high Fourier transform ion cyclotron resonance mass spectrometer (FT-ICR MS). Potential drivers of DOM, including climatic parameters, bacterial community structure and physicochemical factors, were collected. To finely delineate the activity gradient of DOM molecules, we developed a metabolome binning approach to classify DOM molecules into distinct of groups (or bins) based on their potential transformations. The number of molecules in each bin was set as equal, with the first bin having the minimum potential transformations and the last the maximum. The composition and assembly mechanisms of molecular assemblages within each bin and their association with biotic and abiotic variables were explored to test our hypotheses. This molecular level binning approach avoids masking or overlooking information on inactive molecular assemblages that can occur when performing analysis at the compositional level. That is, DOM samples have molecules with the number of putative biochemical transformations ranging from zero to hundreds (Li et al., 2023a), but the mean or intensity-weighted mean value of molecular potential transformations is typically greater than 10 (Hu et al., 2022b; Ryan et al., 2024). As a result, the analysis at the compositional level often masked assembly information of inactive molecules with lower transformations. Collectively, this study has three major objectives: (*i*) determine the composition of DOM in sediments of lake systems; (*ii*) explore the extent to which the composition and assembly of DOM vary with the activity of DOM molecules; and (*iii*) elucidate the biotic and abiotic drivers of DOM composition along the gradient of molecular activity.

## Materials and methods

### Sediment sampling

In total, 63 sediments were collected from freshwater systems during the short period of the 9^th^ of July to the 18^th^ of August 2020. These freshwater systems range from tropical to cold temperate climatic zones in China, with latitudes from 18°48′ to 49°34′LN and longitudes from 102°51′ to 129°36′LE. The mean annual temperature (MAT) of these systems ranges from 2.85 to 25.65°C (Fig. 1a), and mean annual precipitation (MAP) from 175 to 2,362 mm. At each site, we collected the upper 5 cm sediment and the associated surface 20 cm water three times from the lakeshore at the water depth of 0.5 to 1 meter. These samples were mixed to obtain one sediment and one water sample (Wen et al., 2024). All samples were kept in the sterile polyethylene terephthalate bottle, wrapped in ice packs and delivered rapidly to the laboratory for further analysis. Notably, 59 sediments were sampled from ecosystems such as natural lakes and artificial lakes and/or reservoirs, and an additional four sediments were collected from fluvial environments because of limited access to lake sediments at these sites. The fluvial sediments were also included in our analyses as their water depths were similar to the sampled lakes, and their inclusion would not confound the relationships between DOM composition, assembly processes, and potential transformations.

**Fig. 1.**
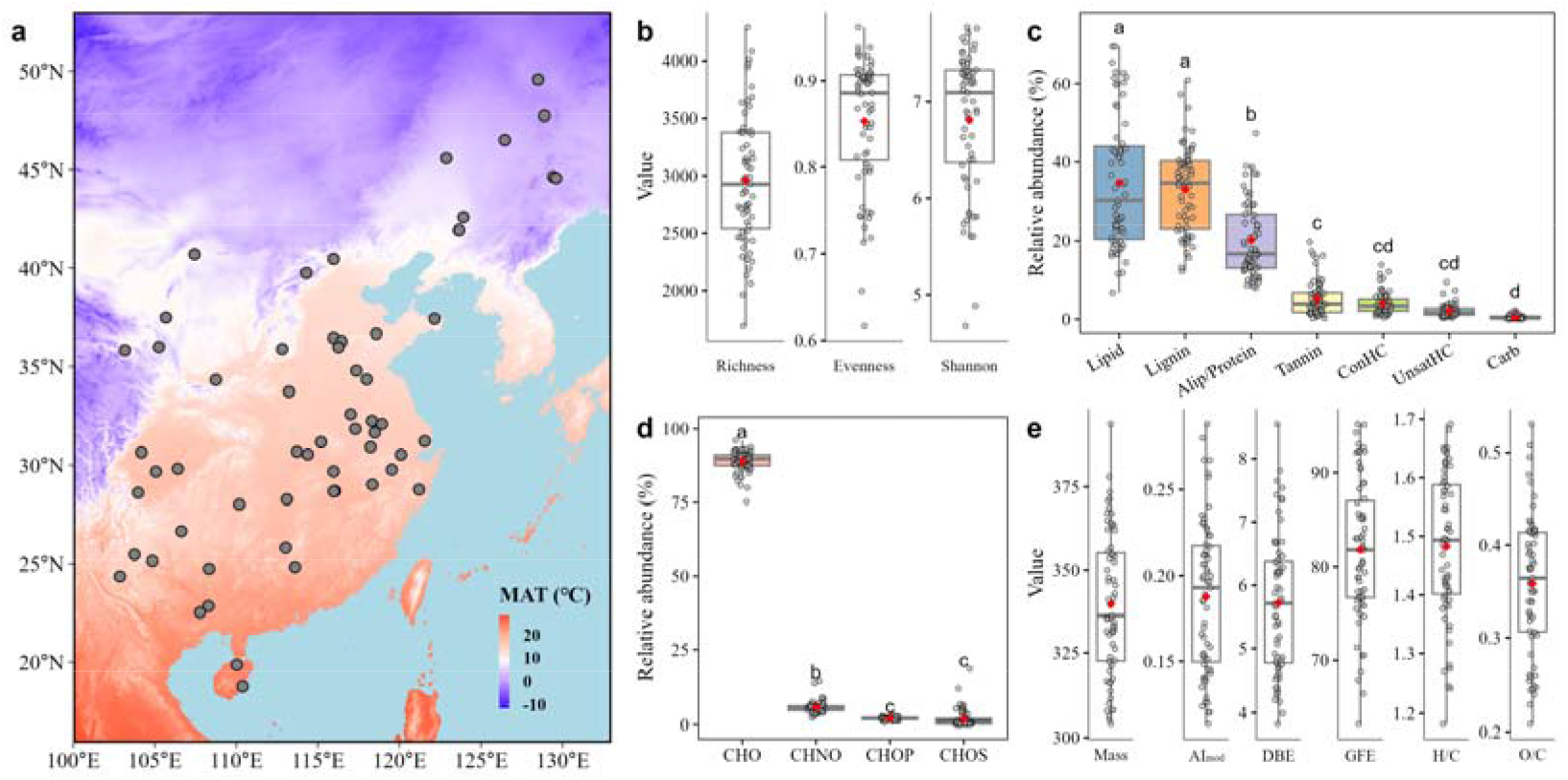
Chemodiversity of sediment DOM in freshwater ecosystems. (a) Sampling map; (b) Alpha diversity of DOM; (c) Relative abundance of DOM compounds; (d) Relative abundance of element combinations of DOM; (e) Intensity-weighted mean values of functional traits of DOM. Lowercase letters in panels (c) and (d) represent group differences determined by LSD test after the ANOVA. Red dots in panels (b-e) represent mean values.

### Climatic, physicochemical and microbial variables

We explored a set of variables potentially associated with the composition and assembly processes of sediment DOM. In brief, climatic variables (i.e., MAT and MAP) were collected using the WorldClim Version 2.1 (Fick and Hijmans, 2017). Physicochemical variables consist of total phosphorus (TP), total nitrogen (TN), electrical conductivity (EC) and pH of water samples, and TP, TN, EC, and pH, total organic carbon (TOC) and dissolved organic carbon (DOC) of sediment samples. TP and TN were quantified using a continuous-flow analyser (Skalar San++, Netherlands). EC and pH were measured directly with a EC (COM-100, HM Digital, USA) and pH probe (SX-610, China), respectively. Sediment TOC was measured with a solid TOC analyzer (SSM- 5000A, Shimadzu, Japan). DOC was analyzed with a Torch TOC analyzer (Teledyne Tekmar, USA). Sediment bacterial communities were profiled via high-throughput 16S rRNA gene sequencing performed on the Illumina MiSeq platform. Data analysis based on amplicon sequence variants (ASVs) was processed using the DADA2 package V1.18 (Callahan et al., 2016). Sequences were rarefied to an equal depth of 32,718 per sample with the “rrarefy” function from the R package “vegan” V2.4.6 (Dixon, 2003). Further details on the bacterial community analyses are provided in our previous study (Wen et al., 2024).

### FT-ICR MS analyses of sediment DOM

First, we extracted DOM from the sediment using ultra-pure water (0.7 g/30 mL) after ultrasonication for 2 hours (Hu et al., 2022a). The supernatant was obtained by centrifugation, filtration and acidification (pH = 2). Second, we extracted the organic matter from the supernatant using pre-activated Oasis HLB cartridges with methanol (ULC-MS grade) and acidic water (pH = 2). Third, DOM in cartridges was eluted into pre-combusted glass vials using methanol, then dried by purging with high purity nitrogen, and finally stored at - 20L for further analysis. DOM was measured by a 15 Tesla FT-ICR MS (Bruker Daltonics, Billerica, MA) in negative ion mode. Detailed test and calibration parameters are available in Wen et al., 2024. Molecular formulae were assigned to mass spectrum peaks according to rigorous elemental combination rules (Hu et al., 2022a; Tolic et al., 2017), with a total of 8,893 formulae were identified in sediment DOM. According to the stoichiometry of molecules, seven classes were categorized as lipid- (H/CL= L1.5–2.0, O/CL=L0–0.3), lignin- (H/CL=L0.7–1.5, O/CL=L0.1–0.67), aliphatic/protein- (Alip/protein; H/CL=L1.5–2.2, O/CL=L0.3–0.67), condensed aromatic- (ConHC; H/CL=L0.2–0.7, O/CL=L0–0.67), tannin- (H/CL=L0.5–1.5, O/CL=L0.67–1.2), carbohydrate- (Carb; H/CL=L1.5–2, O/CL=L0.67–1.2,) and unsaturated hydrocarbon-like (UnsatHC; H/CL=L0.7–1.5, O/CL=L0–0.1) compounds (Kim et al., 2003).

The intensity-weighted traits were calculated to characterize molecule properties such as bioavailability and degree of saturation (Hu et al., 2022a). These traits included molecular mass, modified aromaticity index (AI_mod_), double bond equivalents (DBE), and standard Gibb’s free energy of carbon oxidation (GFE) (Koch and Dittmar, 2015; LaRowe and Van Cappellen, 2011). It also includes stoichiometry ratios, that is O/C, H/C, P/C, S/C, and N/C . High values of AI_mod_ and GFE and lower values of H/C are interpreted as greater recalcitrance of DOM, and high DBE values imply unsaturation (D’Andrilli et al., 2015).

### Molecular activity

Molecular activity, represented by potential biochemical transformations, was inferred from pairwise mass differences amongst molecules within sediment DOM samples. These differences were compared with a common transformation database within 1 ppm of error (Danczak et al., 2020). For instance, a mass difference of 14.01565 between two molecules could presumably indicate a gain or loss of the –CH_2_ group. For each molecule, the potential transformations indicate the count of transformations the molecule involved in, and a larger number of potential transformations implies a higher molecular activity (Hu et al., 2022b; Koch and Dittmar, 2015).

### Statistical analyses

To evaluate the effects of molecular activity on DOM composition and assembly, we used a moving-window approach that allowed us to categorize all molecules equally into bins based on the rank order of molecular potential transformations. Specifically, all molecules were sorted along a gradient of potential transformations, from minimum to maximum. We then used 800 molecules as the window size (i.e., 1–800, 201–1000, …, 8001–8800 and 8094–8893). This approach generated a total of 42 bins, with bins 1 and 42 containing the lowest and highest mean values of potential transformations, i.e., the least active and most active molecules, respectively. This window size provides enough molecules for the analysis of molecule assembly processes within each bin. In addition, the total number of bins enabled to effectively test for potential relationships between molecular activity and chemodiversity and/or the assembly processes at high resolution. To further validate our results, we also performed additional analyses using molecular window sizes of 600 and 1000.

First, the assembly of molecular assemblages in each bin was evaluated using the beta nearest taxon index (βNTI) ecological model (Dini-Andreote et al., 2015). Initially, we multiplied the scaled stepwise transformation distance matrix by the Euclidean distance matrix of 16 molecular traits (Table S1). The above step generated a transformation-weighted molecular characteristics distance matrix, which was used to perform UPGMA hierarchical clustering analysis to construct the transformation-weighted characteristics dendrogram (TWCD) (Danczak et al., 2020). These analyses were performed using the “hclust” function from the R stats V4.3.1 package. Then, βNTI analysis for each bin was performed according to the abundance table. Specifically, the TWCD and a subset of the abundance table for each bin were used to calculate the beta mean nearest taxon distance (βMNTD), which assesses the average phylogenetic distance to the nearest relative between pairs of assemblages (Hu et al., 2022b). The βNTI was calculated as the follows:

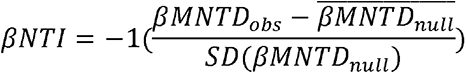

where *βMNTD_obS_* represents the βMNTD of observed molecular assemblages, 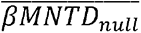 and *SD*(*βMNTD_null_*) represent the average and standard deviation βMNTD of null assemblages, respectively. We performed 999 randomized null assemblages for βNTI analyses, using the “qpen” function from the R package iCAMP V1.5.12 (Ning et al., 2020). Molecular assemblage was interpreted to be structured by deterministic processes when |βNTI| > 2, indicating either variable selection (βNTI > 2) or homogeneous selection (βNTI < -2). If |βNTI| was ≤ 2, the assemblage was interpreted to be structured by stochastic processes. To improve the robustness of our analyses, we also constructed a molecular characteristics dendrogram (MCD) and a transformation-based dendrogram (TD), which were used as inputs for βNTI analyses. The MCD and TD were constructed using 16 molecular traits in Table S1 and molecular potential transformations, respectively. The scripts used to construct these dendrograms can be accessed freely at https://github.com/danczakre/Meta-Metabolome_Ecology.

Second, we employed redundancy analysis (RDA) to quantify the variables associated with variation in molecular assembly along molecular activity. This would reveal the relative contribution of climatic, physicochemical, and bacterial variables on the molecular assemblage in each bin. Climatic variables consisted of MAT and MAP. Physicochemical variables included TOC, DOC, TOC/TN ratio, pH, and EC of sediment samples, and EC of water samples. TN in sediment samples was excluded in these analyses due to its high Pearson’s correlation (*P* < 0.001) with sediment TOC. Likewise, TN, TP, and pH of water samples were excluded as they did not significantly contribute to the variation observed in the molecular assemblages. We performed non-metric multidimensional scaling (NMDS) to analyze variations in bacterial communities according to their Bray-Curtis dissimilarity, and then used first two axes of the NMDS as explanatory variables. For response variables, we used first eight axes of the NMDS according to Bray-Curtis dissimilarities of molecular assemblages. RDA was performed with the “rdacca.hp” function from the R package rdacca.hp V1.1.0 (Lai et al., 2022). Total variance of molecular assemblages explained by environmental and bacterial variables and the individual contributions of each predictor were determined.

Third, to explore variations in potential associations between bacterial taxa and DOM along the molecular activity gradient, we constructed co-occurrence bipartite networks. This was performed using data on bacterial ASVs and molecules subjected to Spearman’s rank correlations. To ensure the reliability of these correlations, only the paired species and molecules that appearing in more than 20 samples were used to generate the network. We retained correlation matrices with significance levels below 0.05 (de Vries et al., 2018). As we focused correlations between bacterial taxa and molecules, bacteria-bacteria and molecule-molecule associations were removed from the final network. Furthermore, we extracted sub-networks based on the molecules within the 42 bins, and calculated topological parameters for each sub-network that include the number of nodes, number of links, average degree, diameter and average path length. We used the “net_properties” function from the R package ggClusterNet V0.1.0 to compute these topological parameters (Wen et al., 2022), and visualized these networks using Gephi (V0.10.1).

We calculated alpha diversity of DOM, including richness, evenness, and Shannon index, as described in Text S1. Relationships between compound abundances and/or DOM traits with molecular activity were examined using linear or quadratic models, depending on the smaller value of Akaike’s information criterion (Akaike, 1974). Regression analysis was used to model the variations in traits and assembly processes of molecular assemblages for each bin along the molecular activity gradient. Least squares algorithms were used to test for the relationships between the relative contribution of potential predictors or network parameters and molecular activity. All of the above statistics were conducted in R (version 4.3.1).

## Results and discussion

### Chemodiversity of sediment DOM

A total of 8,893 formulae were identified in sediment DOM, with the molecular richness ranging from 1,690 to 4,293 (Fig. 1b). The evenness of DOM fell between 0.62 and 0.96, with a mean value of 0.85 that was similar to that observed in three drained wetland soils (Wang et al., 2023). Shannon diversity ranged from 4.68 to 7.78, and was lower than that observed in 45 lake waters (7.23 to 8.38) (Luo et al., 2022) and 11 lakeshore surface sediments (7.01 to 8.2) (Li et al., 2023c). We found DOM to be dominated by lipid- (34.8 ± 16.93%) and lignin- like (33.01 ± 11.01%) compounds, followed by aliphatic/protein-like (20.11 ± 9.4%), while the other compounds such as tannin- or condensed aromatic-like contributed less than 5.28% (Fig. 1c). The dominance of lignin-like compounds was consistent with previous studies in water systems and inland sediments (Li et al., 2023c), and coastal wetlands (Li et al., 2022), which likely attributed to the low bioavailability of these molecules. A distinct finding, however, was the high abundance of lipid-like substance detected in lake sediments. With respect to elemental combinations, DOM was primarily composed of CHO compounds (89.07 ± 3.55%, Fig. 1d). This is in line with the expected DOM composition in inland ecosystems (Luo et al., 2022). Molecules containing heteroatoms such as CHNO, CHOP, and CHOS accounted for 5.8 ± 2%, 2.09 ± 0.52%, and 1.93 ± 3.17%, respectively (Fig. 1d).

The analysis of functional traits of DOM at the compositional level revealed high spatial heterogeneity. The DOM mass ranged from 304 to 393.9 Da (Fig. 1e), which was consistent with the mass range observed in lake waters (Kajan et al., 2023) but lower than that reported for riverine waters that ranged from 518.6 to 567.1 Da (Li et al., 2023b), and for riverine sediments with a mean value of 511 Da (He et al., 2016). The AI_mod_, DBE, GFE, H/C, and O/C values varied over a 2.64, 2.23, 1.5, 1.43 and 2.52-fold range, and changed from 0.11 to 0.29, 3.83 to 8.55, 63.37 to 95.17, 1.18 to 1.69, and 0.21 to 0.53, respectively (Fig. 1e). These trait values exhibited a relatively unique range compared to other habitats, such as smaller AI_mod_ and DBE, and greater H/C compared to riverine sediments (Han et al., 2024). This high functional diversity of traits of DOM in lake sediments is partly attributed to variations in climatic and anthropogenic variables at large spatial scales that structure DOM chemodiversity (Kida et al., 2023; Lehmann et al., 2020).

### Varying sediment DOM chemodiversity with molecular activity

Of all 8,893 unique molecules detected in sediment DOM, the putative biochemical transformations ranged from 0 to 79. Towards high potential transformations, molecular frequency showed a decreasing trend (Fig. 2a), while the molecular relative abundances increased significantly (*P* < 0.001, Fig. 2b). Furthermore, we found a decreasing beta-diversity along the potential transformations, as evidenced by the decrease in Bray-Curtis dissimilarities of molecular assemblages across bins (R^2^ = 0.65, *P* < 0.001, Fig. 2c). Specifically, toward bins of high potential transformations, molecular counts of lignin- and tannin-like compounds within bins increased significantly, with R^2^ of 0.98 and 0.58, respectively (*P* < 0.001, Figs. 2d, S1). Conversely, the counts of aliphatic/protein-, condensed aromatic-, unsaturated hydrocarbon- and carbohydrate-like compounds decreased significantly and reached zero, with R^2^_adj_ values ranging from 0.46 to 0.91. The count of lipid-like compounds, however, showed a hump-shaped pattern (R^2^_adj_ = 0.66, Figs. 2d, S1). Regarding the relative abundance of compound classes, lipid-like substances exhibited a hump-shaped pattern, which was opposite to lignin-like compounds that showed a U-shaped pattern (Figs. 2e, S2). As for condensed aromatic-, unsaturated hydrocarbon- and carbohydrate-like compounds, each of their relative abundances decreased significantly (*P* < 0.001), in line with the variation in molecular counts within the bins (Fig. S2). Taken together, these results demonstrate that molecular assemblages in bins of high molecular activity are increasingly dominated by universal, recalcitrant lignin-like compounds (Freeman et al., 2024), and thus are more homogeneous.

**Fig. 2.**
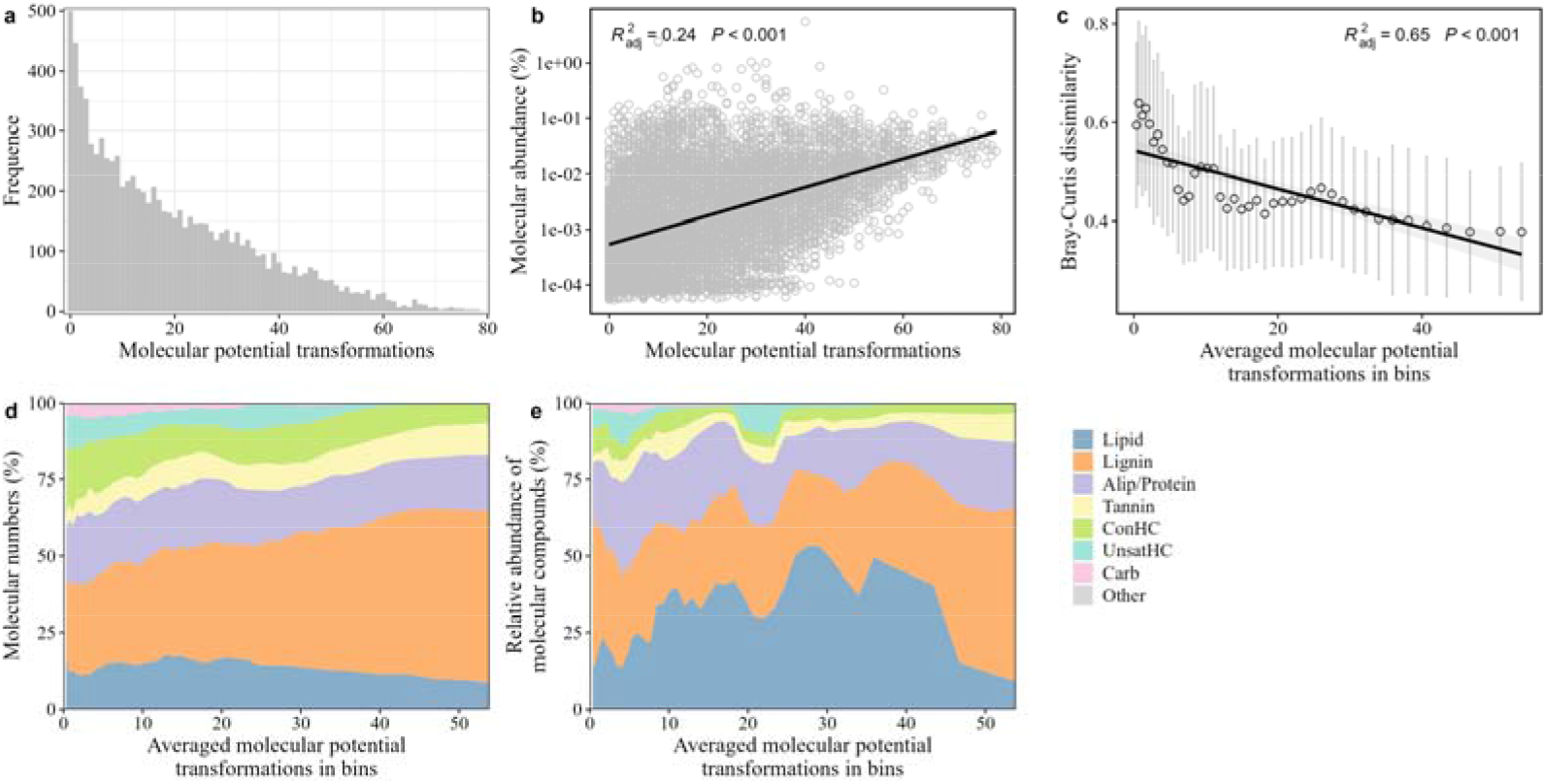
Variations in chemodiversity of sediment DOM along the gradient of molecular activity represented by potential transformations. Panels (a) and (b) depict variations in the frequency and relative abundance of molecules in DOM in relation to their potential transformations; (c) Bray-Curtis dissimilarity of molecular assemblages as a function of the averaged potential transformations within bins, with dots representing means and error bars representing standard deviations; Panels (d) and (e) display variations in molecular numbers and relative abundance of different classes within bins along the gradient of potential transformations. Bins: A total of 42 bins were generated along the gradient of molecular potential transformations, with each bin containing 800 molecules, bin1 having the lowest value and bin42 having the highest value of molecular potential transformation. Alip/Protein: aliphatic/protein- like compounds. ConHC: condensed aromatic-like compounds. UnsarHC: unsaturated hydrocarbon-like compounds. Carb: carbohydrate-like compounds.

We found variations in DOM traits along the gradient of potential transformations to exhibit complex and distinct patterns (Fig. S3). These trait values were calculated by the intensity-weighted means and further averaged across all samples within bins. The molecular mass was found to decrease rapidly from 482 Da to 324 Da as potential transformations increased from 0 to 10, remaining stable thereafter (Fig. S3a). The AI_mod_ and DBE values were found to sharply decrease when potential transformations increased from 0 to 16, later showing an upward trend, whereas H/C displayed an opposite pattern (Figs. S3b, c, f). Values of GFE and O/C had opposite patterns along the transformation gradient, with extreme values observed at the transformation value of 29 (Figs. S3d, e). For features characterizing heteroatoms, such as N/C, P/C, and S/C, it was found that they decrease and converge to zero with increasing potential transformations (Figs. S3g-i). This indicates that active compounds are less likely to contain heteroatoms.

### Sediment DOM assembly integrated with molecular activity

For the assembly of DOM, we found that the interplay of stochastic and deterministic processes varies across molecular activity. Specifically, the relative contribution of stochastic processes sharply dropped from 93% to 5.1% as potential transformations increased from 0.4 to 14, having a slight increase or being relatively stable thereafter (Fig. 3). Conversely, the contribution of variable selection improved from 6.2% to 95%, and remained relatively stable thereafter. The contribution of homogeneous selection was found to be < 1.6% over the full range of potential transformations. The robustness of these findings was corroborated by similar analyses applied to MCD or TD, and based on TWCD with window sizes of 600 and 1,000 molecules (Fig. S4). Our findings align with previous studies showing that biochemically active molecules (with higher transformations) are mostly structured by deterministic processes (Danczak et al., 2020). We quantitatively characterized the variation of DOM assembly processes along the gradient of potential transformations at the molecular level. This approach contributes to the understanding of the mechanisms structuring the molecular assemblages within specific segments of potential transformations at the compositional level. For instance, we observed DOM at the compositional level to be structured by deterministic processes with a contribution of 99.6% (Fig. S5). The mean or intensity-weighted mean values of potential transformations for the 63 DOM samples ranged from 23.49 to 29.30 and 21.93 to 30.98, respectively. These results suggest that DOM is highly active at the compositional level, and could potentially impair the detection of the assembly of inactive molecules. Therefore, we proposed that analysis at the molecular level to be more advantageous to determine the influence of molecular properties on DOM assembly mechanisms (Li et al., 2023a).

**Fig. 3.**
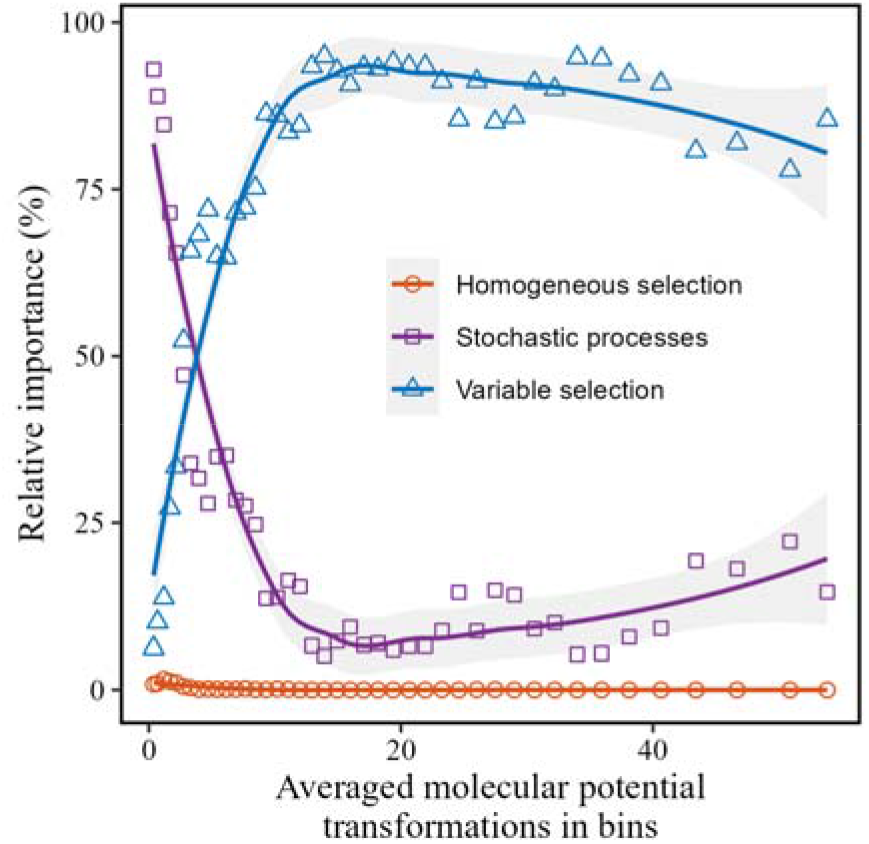
Relative importance of assembly processes structuring the molecular assemblages within bins along the gradient of molecular activity. The molecular activity was represented by potential transformations. Shadings represent 95% confidence intervals.

Variations in the assembly processes of molecular assemblages were also well- predicted by other functional traits. For instance, we found the relative importance of variable selection to decrease monotonically with mass and DBE (*P* < 0.001), with R^2^_adj_ of 0.89 and 0.82, respectively (Figs. 4a, c). Even though the relative importance of variable selection had significant correlations (*P* < 0.01) with traits of AI_mod_, GFE, and H/C, the R^2^_adj_ values were ≤ 0.23 (Figs. 4b, d, f). Furthermore, the molecular assemblages in bins of high transformations displayed high variability in AI_mod_, GFE, O/C, and H/C, but were found to converge at the low molecular mass and DBE (Fig. 4). Thus, we considered mass and DBE as potentially important traits associated with molecular activity to determine the assembly processes of DOM (Li et al., 2023a). Molecular mass and DBE are important properties governing the selective adsorption of DOM by minerals (Lv et al., 2016). Additionally, molecular mass is critical to the photo- and bio-degradation processes of DOM (Catalán et al., 2017; Xu et al., 2022). Molecules with low mass are known to exhibit high photo-reactivity as indicated by high quantum yields (Zhang et al., 2022). These molecules are also preferentially metabolized by bacteria due to their greater labile nature and bioavailability (Roth et al., 2019). The above factors likely contribute to the higher potential transformations of low molecular mass fractions (Fig. S3a). Consequently, biochemical transformations of molecules with changes in traits such as molecular mass and DBE regulate the turnover and composition of DOM.

**Fig. 4.**
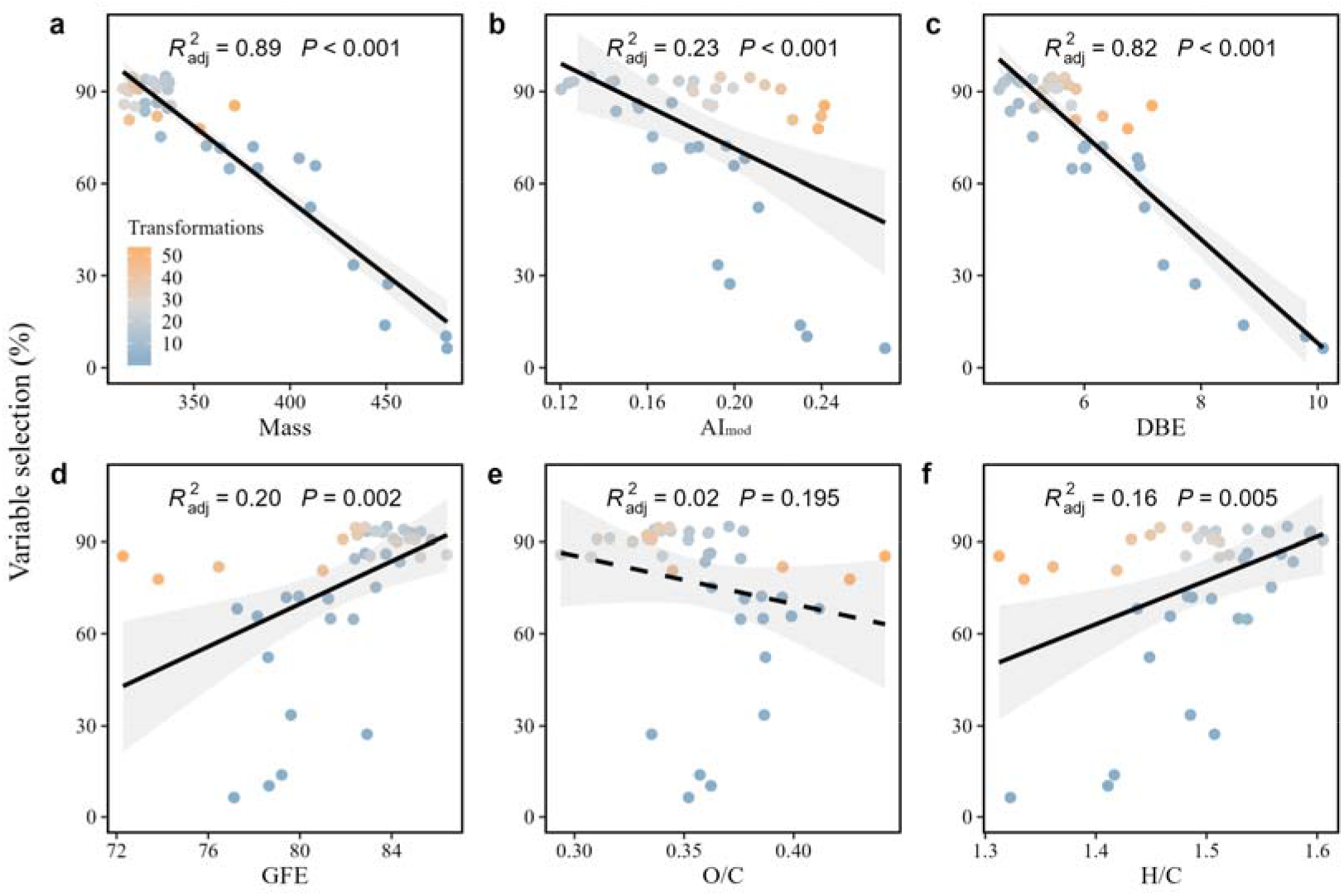
Relationships between the relative importance of variable selection and functional traits of sediment DOM. Lines and shadings indicate linear regression and 95% CI. The colours of the dots in the panels represent the averaged molecular potential transformations in bins.

We found the molecular assemblages to be mostly associated with sediment physicochemical properties, while contributions of climatic and bacterial variables increased with increasing molecular activity. Specifically, the total variance (R^2^_adj_) of molecular assemblages within bins explained by these variables ranged from 16.5% to 24.9% (Fig. 5a), with greater contributions of physicochemical variables (i.e., sediment TOC, EC, and pH) (Figs. 5b, S6). There was an insignificant linear relationship between the total variance explained and potential transformations (Fig. 5a), which stands in contrast with the positive relationship between the importance of deterministic processes and potential transformations (Fig. 3). This could be explained by the fact that sediment DOM composition is dynamically regulated by several drivers, such as benthic fauna, plant composition, and fungi (Taube et al., 2018; Wetzel and Søndergaard, 1998), which were not included as variables in our study. Moreover, the relative contributions of climate (12.99–28.28%) and bacterial composition (11.93–27.6%) were found to increase in bins of high potential transformations, while the relative contribution of physicochemical variables exhibited a decreasing trend (Fig. 5b). The increased effect of bacterial communities was further supported by the increasing complexity of DOM-bacteria bipartite networks (Fig. 6a). In brief, the number of nodes, links, and average degree of networks were found to increase, while the average path length and diameter decreased significantly as the potential transformations of molecular assemblages increased (*P* < 0.001; Fig. 6). In addition, we found that the increase in the number of nodes was mostly attributed to DOM molecules (especially lignin-like compounds, Figs. 6b, S7), while the increase in average degree was mostly attributed to bacterial taxa (especially *Chloroflexi*, *Bacteroidota,* and *Actinobacteriota*, Figs. 6d, S8, S9). These findings imply that the strengthening of network relationships was primarily due to the increase in the number of molecules associated with bacteria, that is, high generality of bacteria (Hu et al., 2022a). Therefore, bacterial taxa in the network may be comparable across molecular assemblages with differing potential transformations (Fig. S7b), but the frequency of bacterial-molecular interactions is likely to vary. Last, molecular assemblages with lower potential transformations were found to display fewer interactions with bacterial taxa, whereas a significant proportion of assemblages in bins of high potential transformations were directly associated with bacteria.

**Fig. 5.**
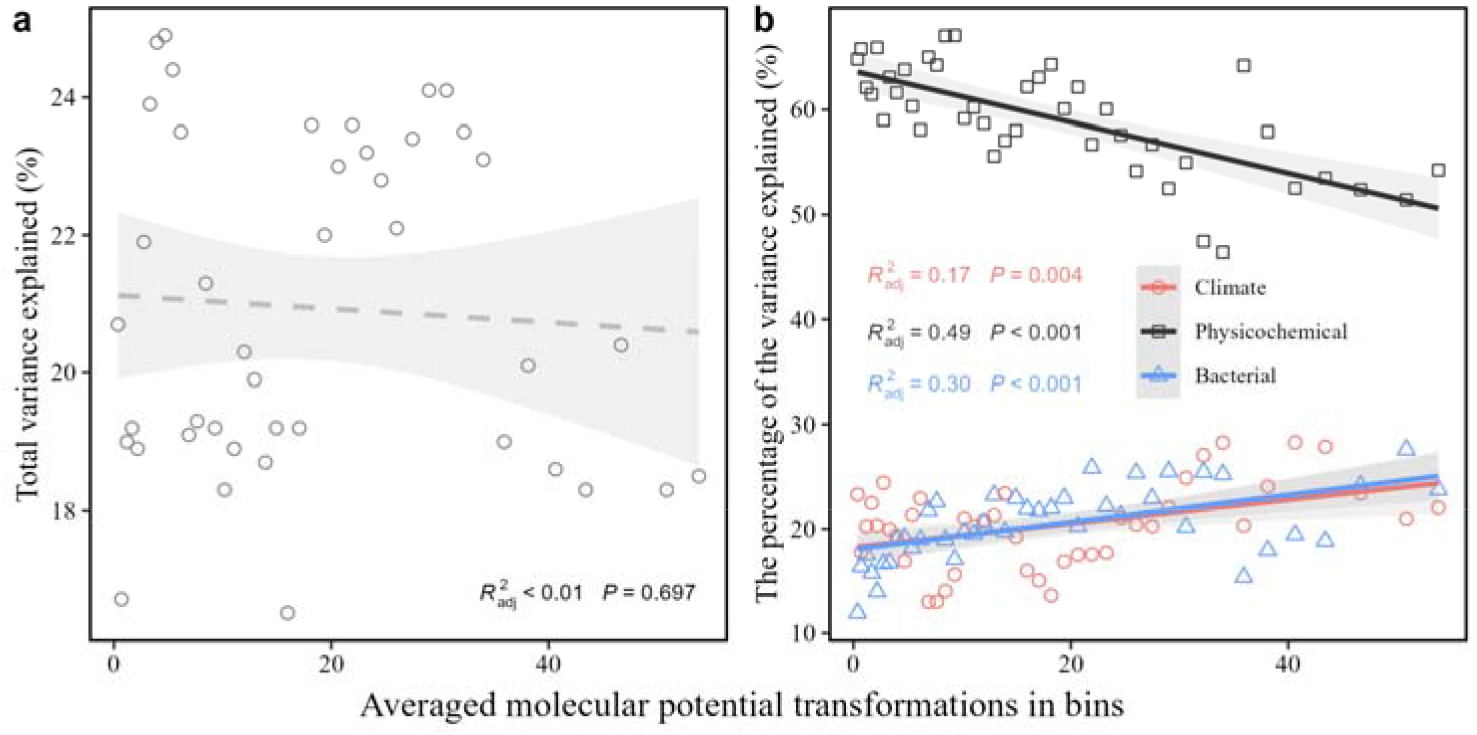
Variances of DOM composition explained by climatic, physicochemical and bacterial community variables along the gradient of molecular activity represented by potential transformations. Lines and shadings indicate linear regression and 95% CI. Climatic variables include mean annual temperature and mean annual precipitation. Physicochemical variables include electrical conductivity of water, and carbon to nitrogen ratio, electrical conductivity, pH, dissolved organic carbon and total organic carbon of sediment.

**Fig. 6.**
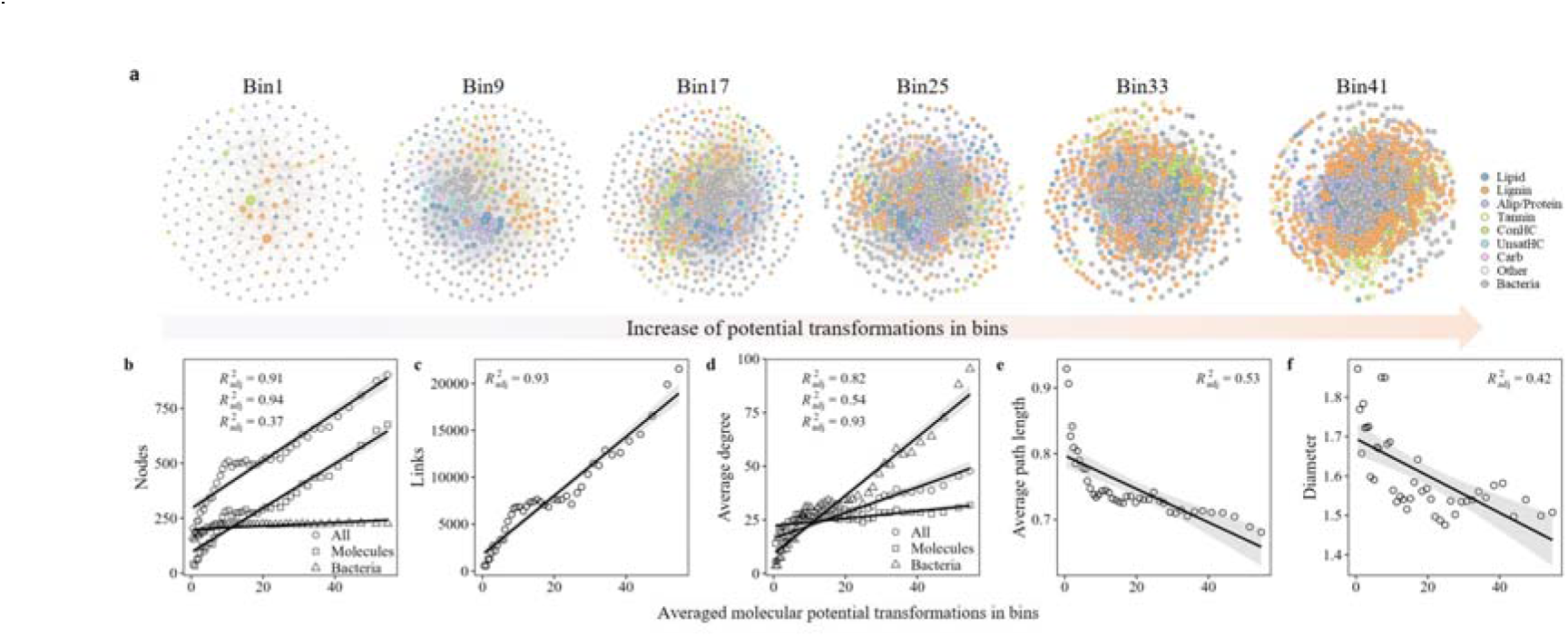
Bacteria-molecules bipartite networks along the gradient of molecular activity represented by potential transformations. (a) visualization of networks; (b-f) Topology properties of networks in relation to potential transformations, including the network number of nodes (b), number of links (c), average degree (d), average path length (e), and diameter (f). Lines and shadings represent linear regression (*P* < 0.001) and 95% confidence intervals. Bins: A total of 42 bins were generated along the gradient of molecular potential transformations, with each bin containing 800 molecules, bin1 having the lowest value and bin42 having the highest value of molecular potential transformation. Alip/Protein: aliphatic/protein-like compounds. ConHC: condensed aromatic-like compounds. UnsarHC: unsaturated hydrocarbon-like compounds. Carb: carbohydrate-like compounds.

## Conclusions

Our large-scale spatial study provides evidence for a strong coupling between the intrinsic activity of DOM molecules and their composition and assembly in lake sediments. In brief, DOM molecules with higher activity tend to be enriched in recalcitrant lignin-like compounds, and exhibit greater compositional similarity across diverse climatic zones. As the potential transformations increased from 0.4 to 14, DOM assembly processes shifted from stochastically to deterministically dominated, indicated that active molecules tend to be assemble deterministically. While physicochemical factors emerged as the primary regulators of DOM composition, the influence of climatic and bacterial variables increased with increasing molecular activity, consistent with the more complex network relationships between the active molecules and bacterial communities. The leveraging of a metabolome binning approach at the molecular level provides the necessary tool to explore the processes and mechanisms structuring DOM that would be obscured via compositional-level analyses. In summary, our findings elucidate the composition, assembly processes and structuring mechanisms of DOM in lake systems. This represents an important step toward advancing our understanding of DOM dynamics in freshwaters and provides new insights into the global carbon cycle.

## Author Contributions

J.W. conceptualized the study. L.H., S.J., A.H. and J.W. collected the samples. K.S.J. analyzed the DOM with the FT-ICR MS. S.W. conducted the statistical analyses and wrote the first draft with the guidance of J.W. S.W. revised the manuscript based on suggestions from J.W., F.D. and A.H. All authors contributed to the intellectual development of this study.

## Notes

The authors declare no competing financial interest.

## Acknowledgments

We appreciate Fanfan Meng, Jinfu Liu, Minglei Ren, Hao Wu, Yanan Zhou, Jianing Xu, Jiyi Wang, Pubo Chen, Huilin Liu, Zhenghua Liu, Weizhen Zhang, Xu Ma, Chaoxun Guo and Yunfeng Wang for their assistance with field sampling. This work was financially supported by research grants from National Natural Science Foundation of China (42225708, 92251304, 42377122, 42307323), Second Tibetan Plateau Scientific Expedition and Research Program (STEP, 2019QZKK0503), Key Laboratory of Lake and Watershed Science for Water Security (NKL2023-QN04), and Science and Technology Planning Project of NIGLAS (NIGLAS2022GS09).

## Notes

### Competing Interest Statement

The authors have declared no competing interest.

