## Supplementary material for "Molecular activity mediates the composition and assembly of dissolved organic matter in lake sediments": Suppleemental Information

**This file includes:**

Supplementary Table (S1)

Supplementary Text (S1)

Supplementary Figures (S1–S9)

**Supplementary Table**

**Table S1.** Molecular traits used to construct the molecular characteristics dendrogram and the transformation-weighted characteristics dendrogram.

| Molecular traits | Description |
| --- | --- |
| Mass | Mass of DOM molecules |
| C | Number of C atoms in DOM molecules |
| AI_mod_ | Modified aromacity index of DOM molecules |
| DBE | Double bond equivalents of DOM molecules |
| DBE-O | DBE minus number of oxygens |
| DBE-AI | DBE minus AI |
| GFE | Standard Gibb’s Free Energy of carbon oxidation of DOM molecules |
| Kdefect_CH2_ | Kendrick Defect of DOM molecules |
| NOSC | Nominal oxidation state of carbon of DOM molecules |
| O/C | The ratio of O number to C number in DOM molecules |
| H/C | The ratio of H number to C number in DOM molecules |
| N/C | The ratio of N number to C number in DOM molecules |
| P/C | The ratio of P number to C number in DOM molecules |
| N/P | The ratio of N number to P number in DOM molecules |
| S/C | The ratio of S number to C number in DOM molecules |
| Y_met_ | Carbon use efficient |

**Text S1**

We calculated the Richness (S), Shannon diversity (H’), and Evenness (E) of sediment DOM. Richness was calculated as the number of molecular species. Shannon diversity and Evenness were calculated as follows (Dixon, 2003):

$$H’=-\sum P_{i}\ln(P_{i})$$

(1)

$$E=H’/\ln(S)$$

(2)

Where $P_{i}$ is the relative abundance of molecules $i$.

**Supplementary Figures**

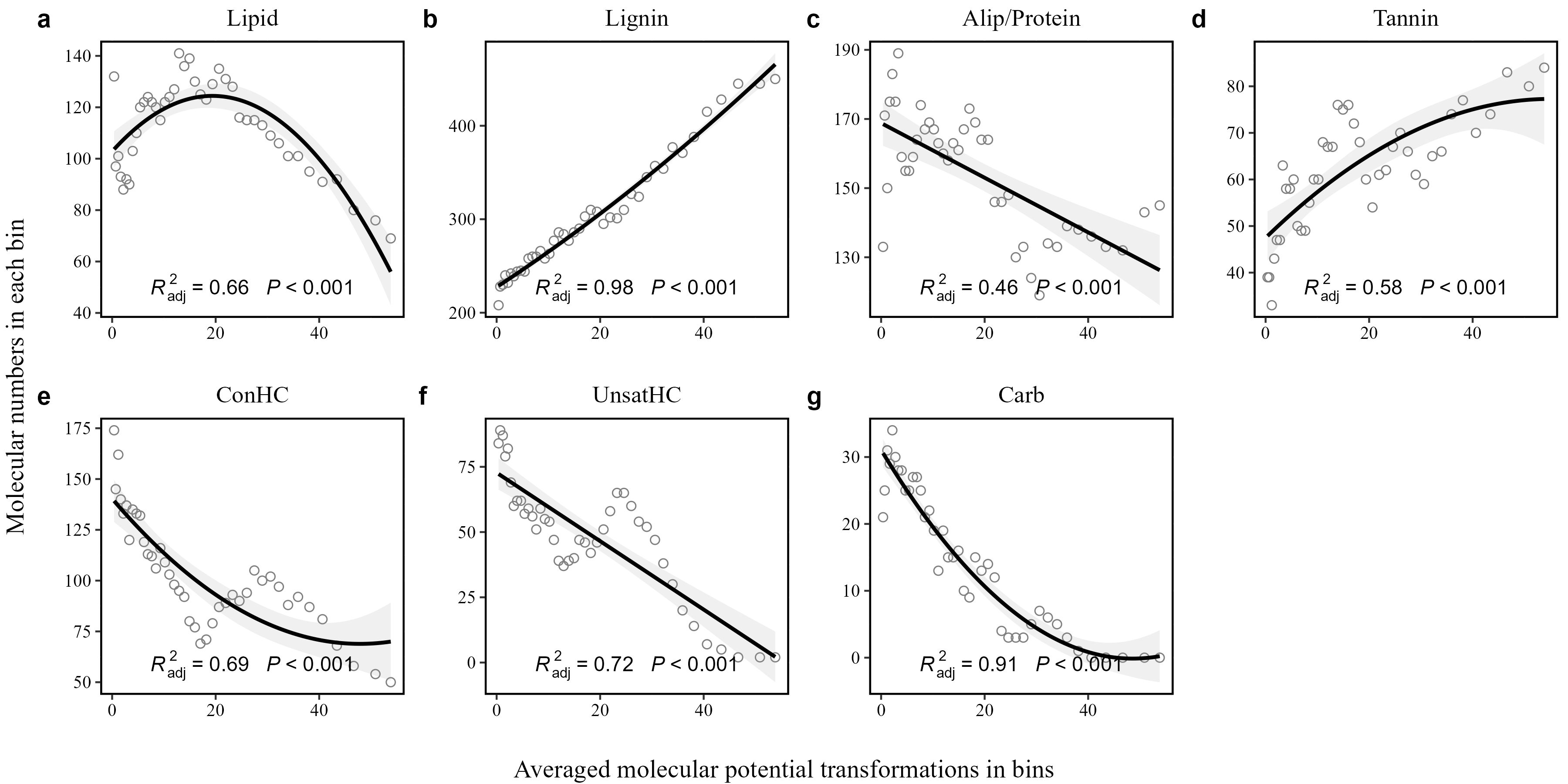

Figure S1. The relationships between the molecular numbers and averaged potential transformations within bins. Bins: A total of 42 bins were generated along the gradient of molecular potential transformations, with each bin containing 800 molecules, bin1 having the lowest value and bin42 having the highest value of molecular potential transformation. Alip/Protein: aliphatic/protein-like compounds. ConHC: condensed aromatic-like compounds. UnsarHC: unsaturated hydrocarbon-like compounds. Carb: carbohydrate-like compounds.

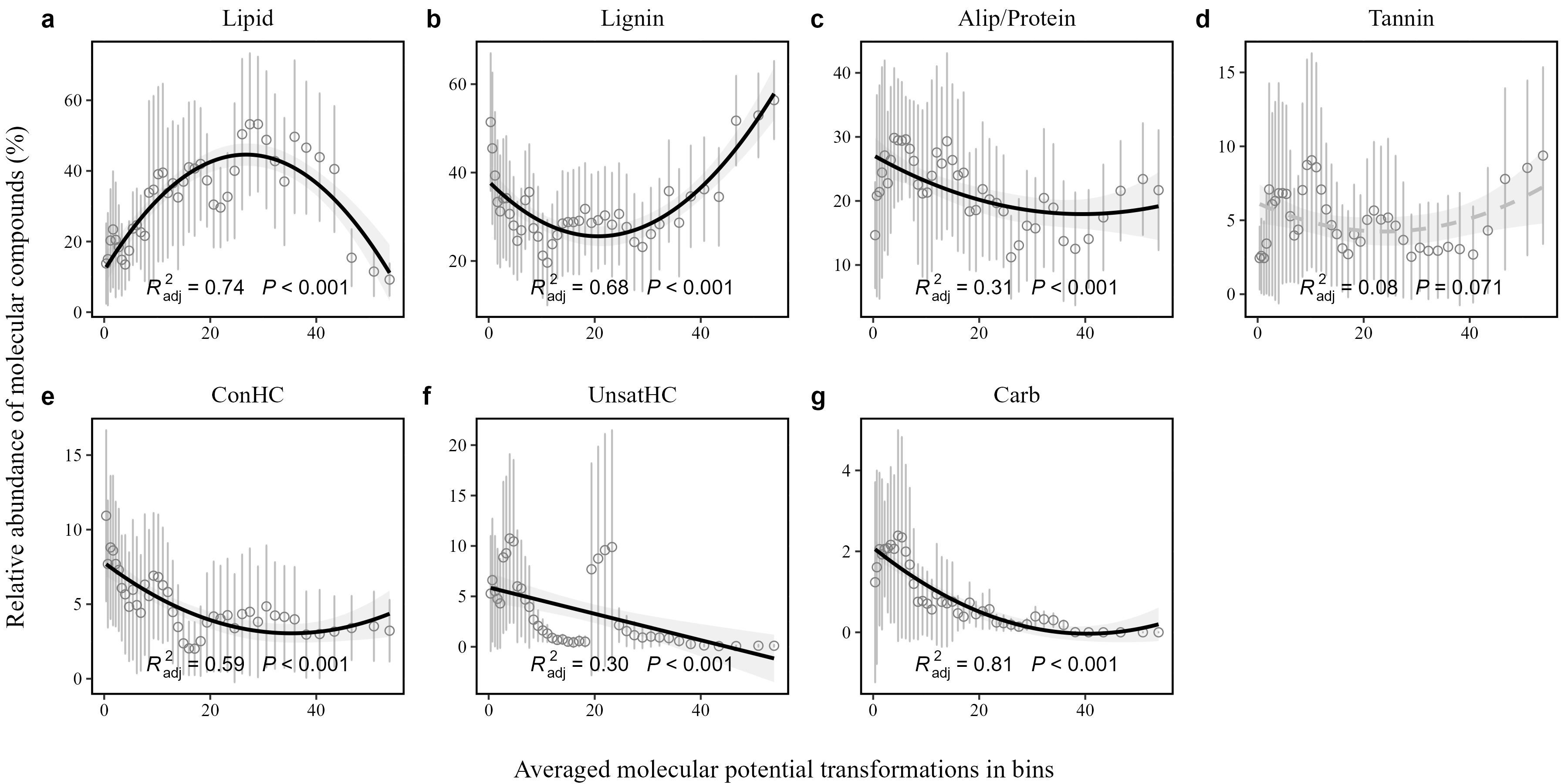

Figure S2. The relationships between the relative abundance of each chemical compound and averaged potential transformations within bins. The dots represent mean values and error bars represent standard deviations. Shadings represent 95% CI. Bins: A total of 42 bins were generated along the gradient of molecular potential transformations, with each bin containing 800 molecules, bin1 having the lowest value and bin42 having the highest value of molecular potential transformation.

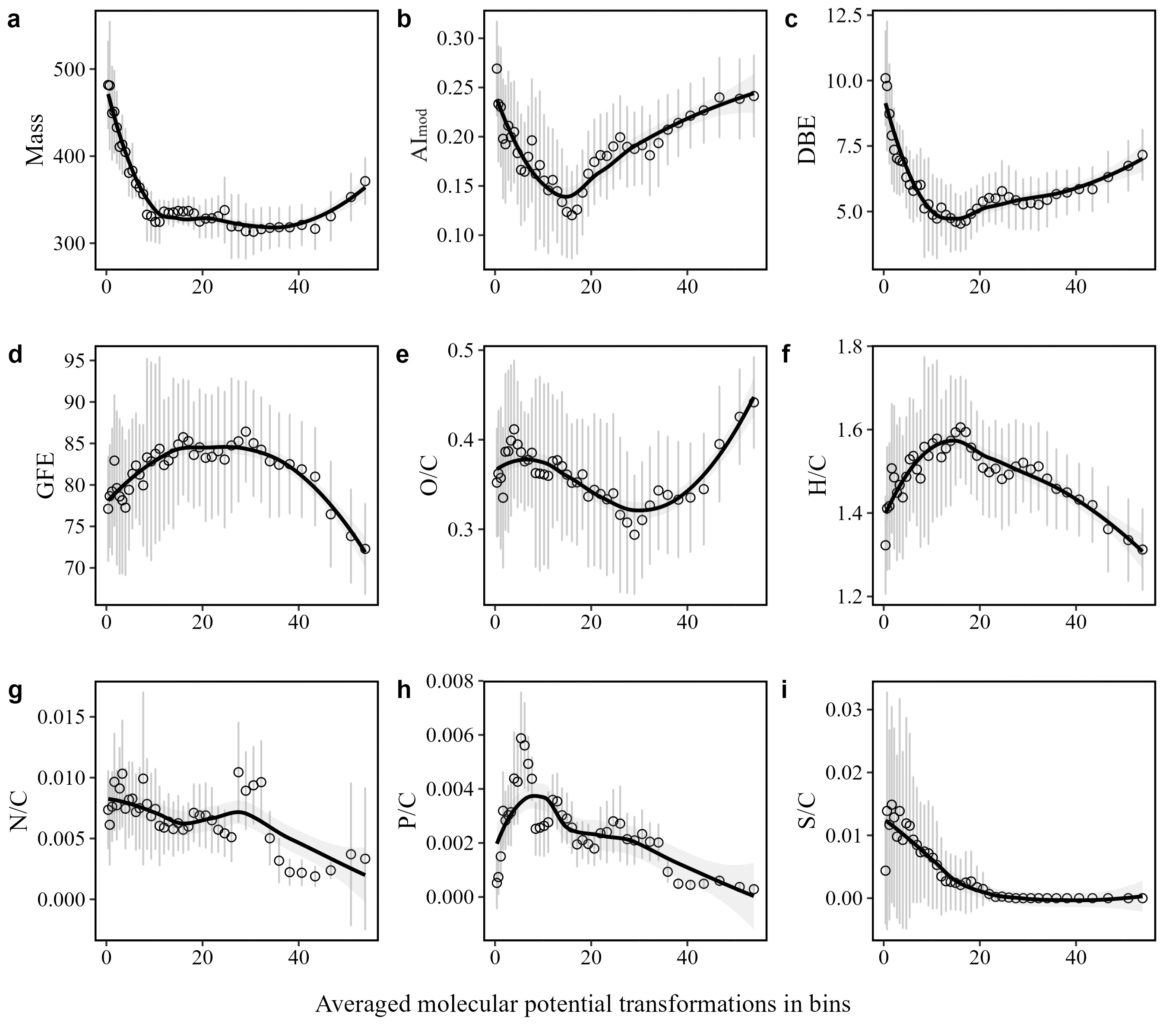

Figure S3. Intensity weighted mean values of traits of molecular assemblages in relation to averaged potential transformations within bins. The dots represent mean values and error bars represent standard deviations. Shadings represent 95% CI.

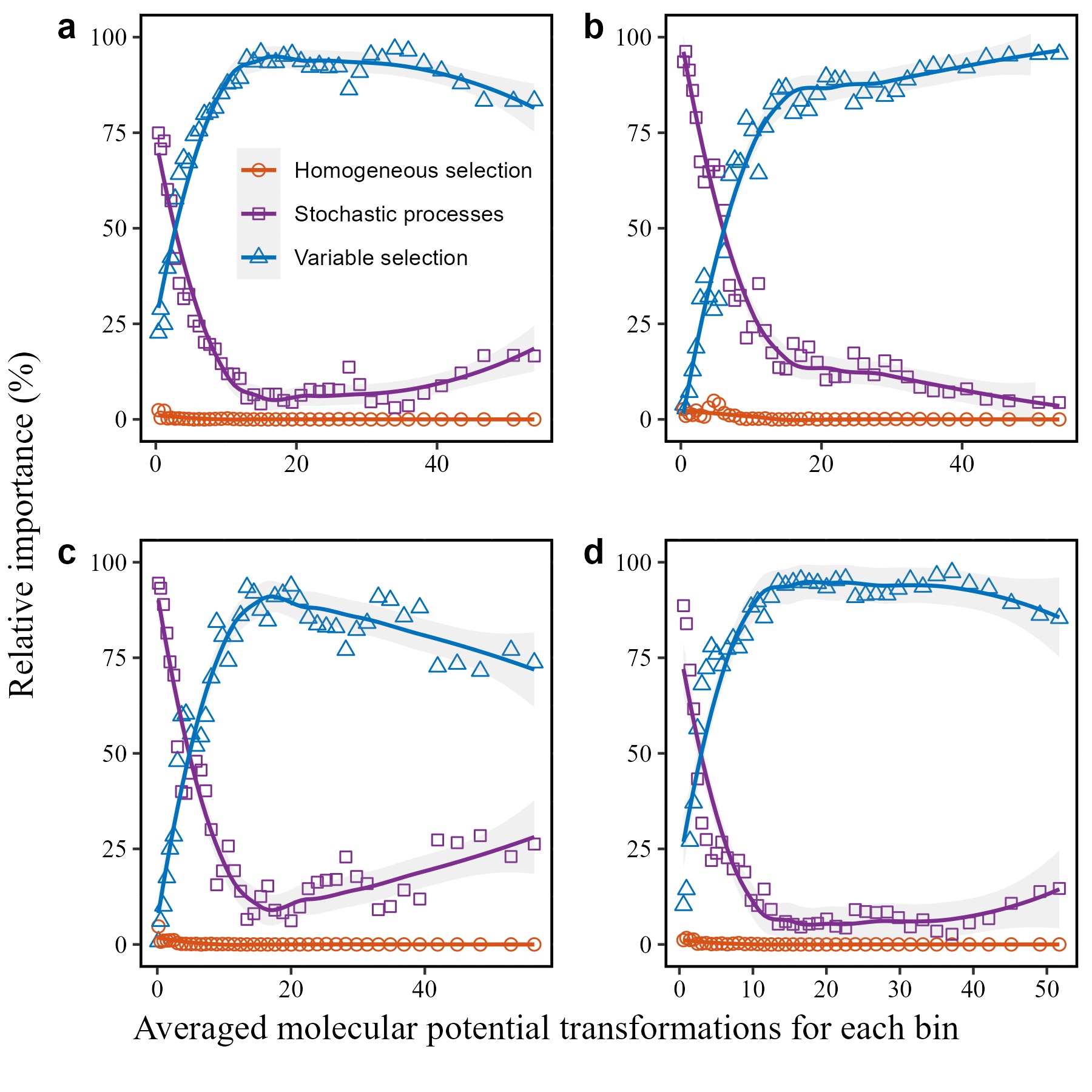

Figure S4. Relative importance of assembly processes structuring DOM composition in sediments along with the potential transformations. (a) Assembly processes calculated using the molecular characteristics dendrogram with each bin containing 800 molecules; (b) Assembly processes calculated using the transformation-based dendrogram with each bin containing 800 molecules; (c) Assembly processes calculated using the transformation-weighted characteristics dendrogram (TWCD) with each bin containing 600 molecules; (d) Assembly processes calculated using the TWCD with each bin containing 1000 molecules. Shadings represent 95% CI.

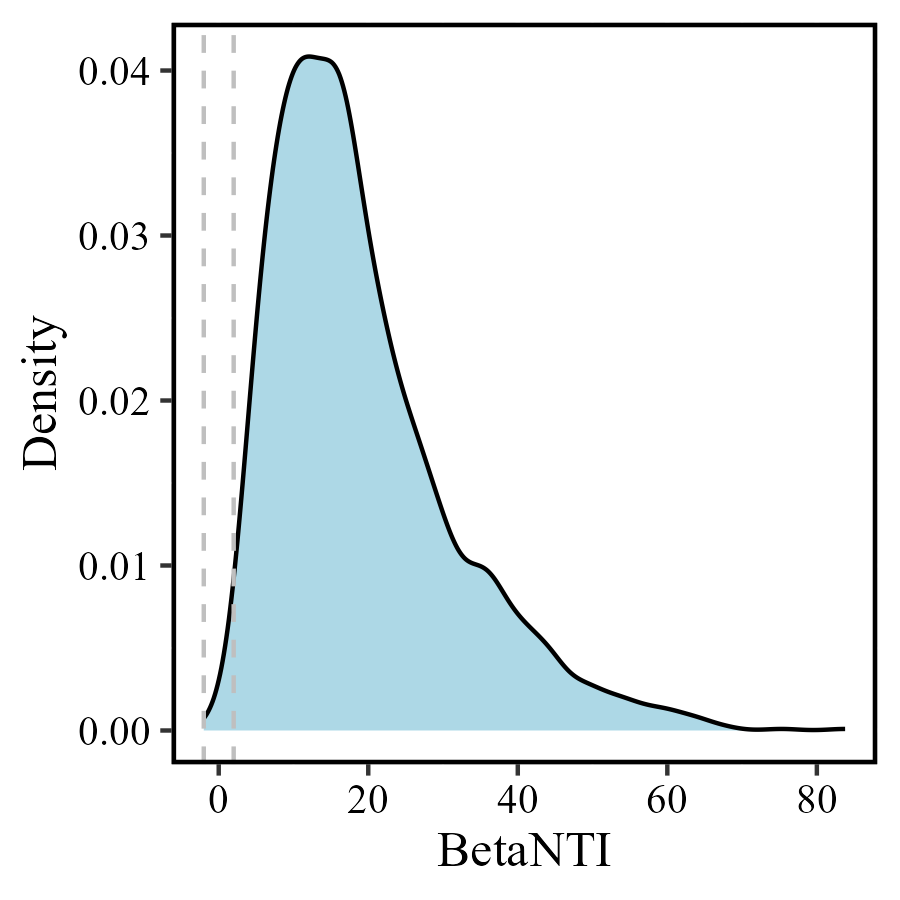

Figure S5. Density plot of βNTI based on TWCD for sediment DOM at the compositional level. Dashed lines indicate the assembly processes threshold of |βNTI| < 2. βNTI: beta nearest taxon index.

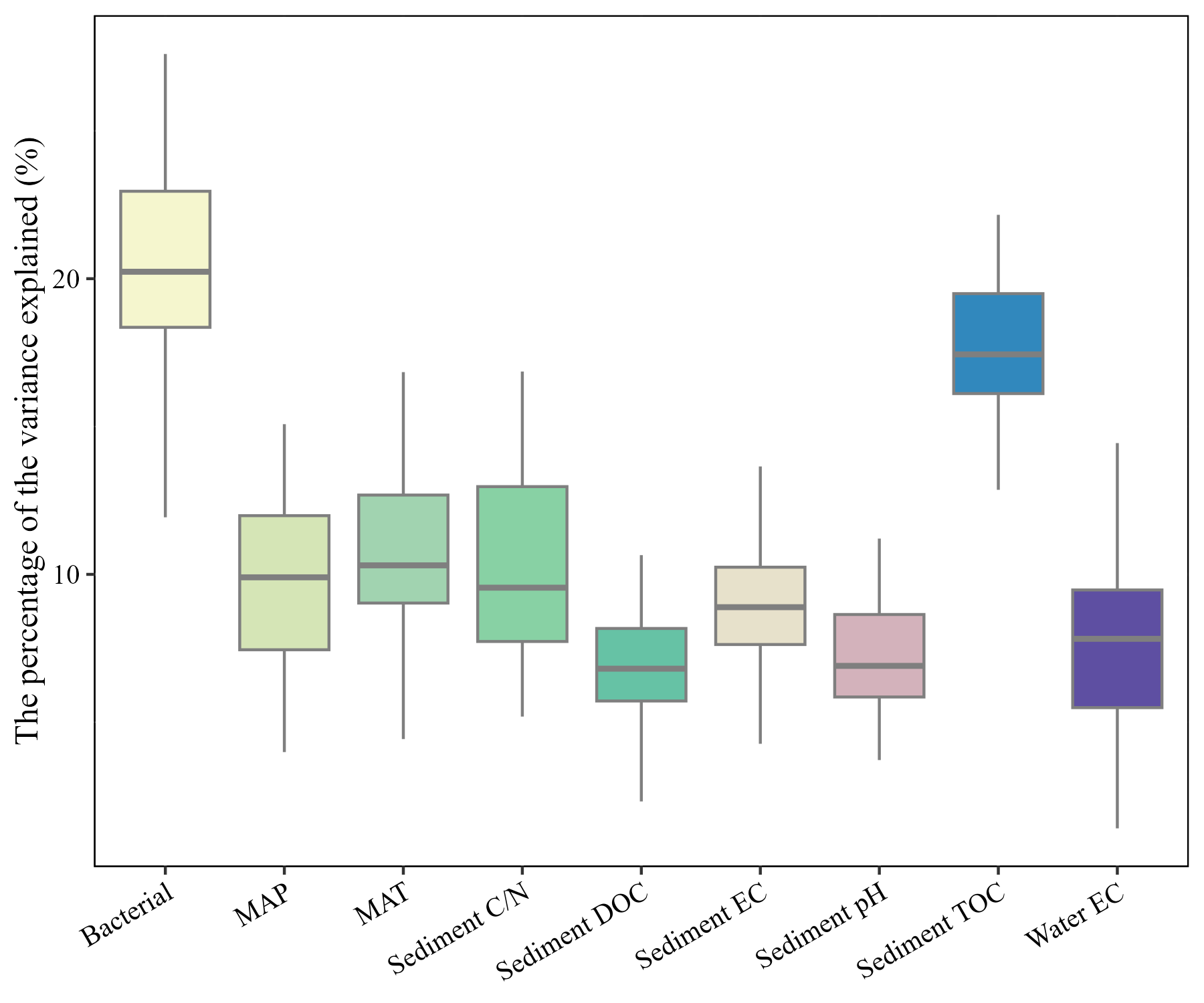

Figure S6. Percentage variance in DOM explained by distinct variables. MAT: mean annual temperature. MAP: mean annual precipitation. C/N: carbon to nitrogen ratio. DOC: dissolved organic carbon. EC: electrical conductivity. TOC: total organic carbon.

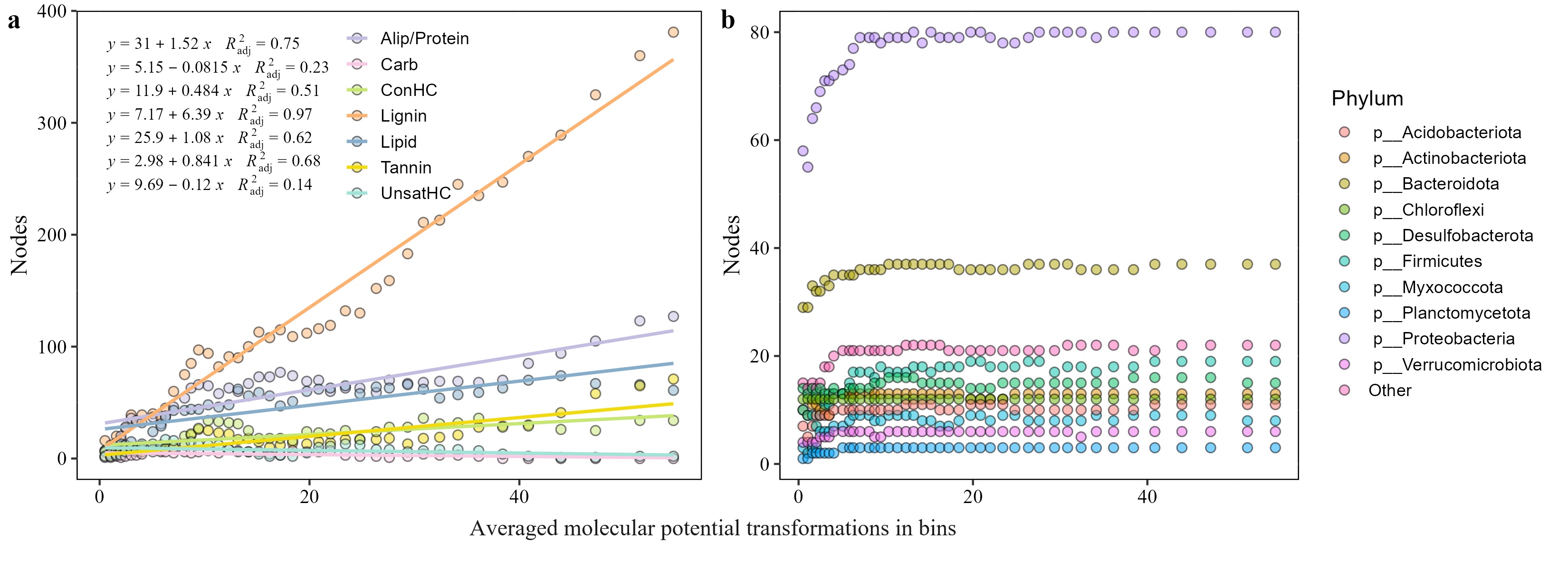

Figure S7. Molecular classes (a) and bacterial phyla (b) present as network nodes in relation to potential transformations.

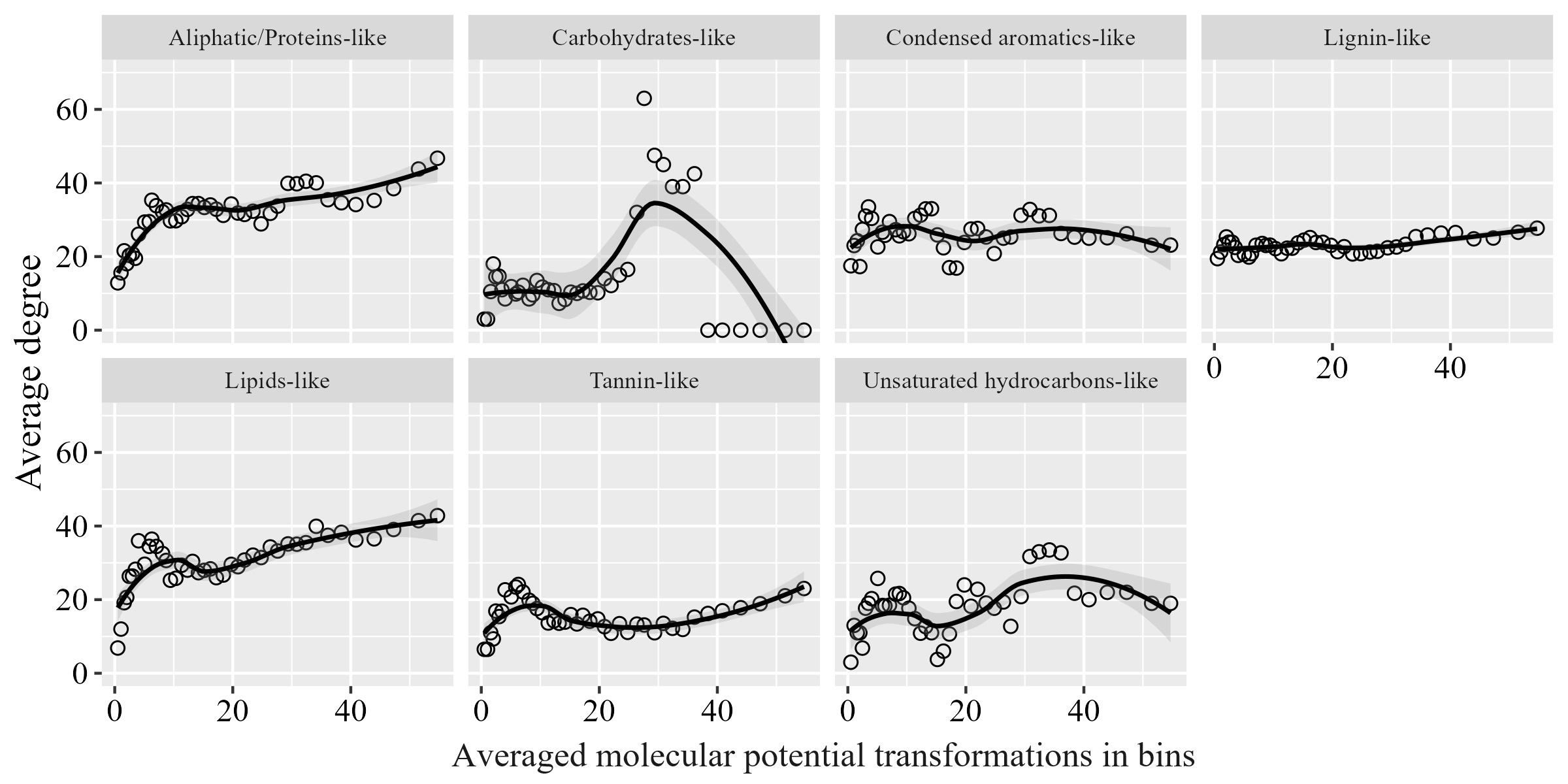

Figure S8. Relationships between the average degree of different classes of molecules and potential transformations. Shadings represent 95% CI.

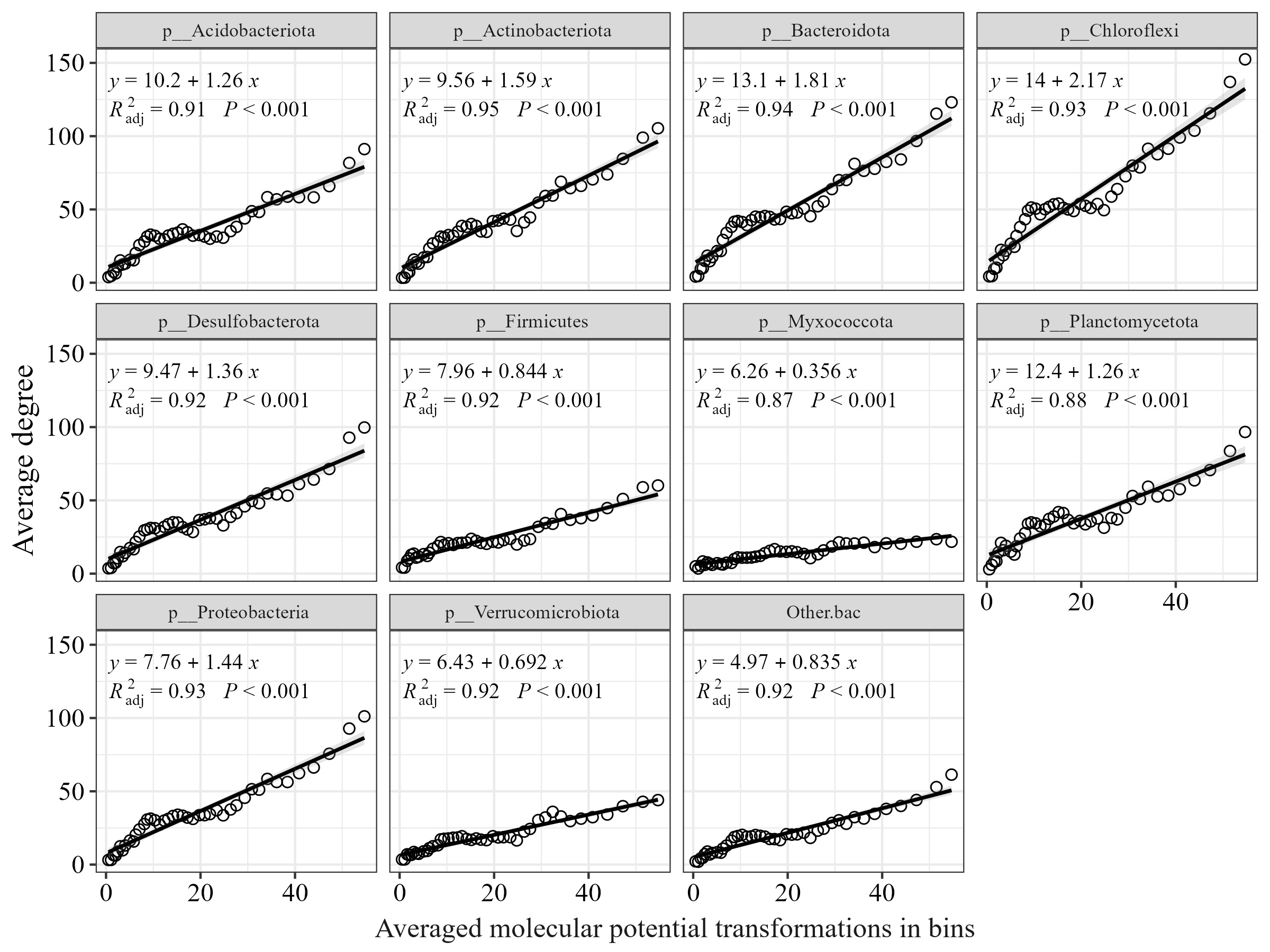

Figure S9. Relationships between the average degree of bacteria phyla and potential transformations. Lines and shadings represent linear regression and 95% CI.
